# Latent Feature Representations for Human Gene Expression Data Improve Phenotypic Predictions

**DOI:** 10.1101/2020.10.15.340802

**Authors:** Yannis Pantazis, Christos Tselas, Kleanthi Lakiotaki, Vincenzo Lagani, Ioannis Tsamardinos

## Abstract

High-throughput technologies such as microarrays and RNA-sequencing (RNA-seq) allow to precisely quantify transcriptomic profiles, generating datasets that are inevitably high-dimensional. In this work, we investigate whether the whole human transcriptome can be represented in a compressed, low dimensional latent space without loosing relevant information. We thus constructed low-dimensional latent feature spaces of the human genome, by utilizing three dimensionality reduction approaches and a diverse set of curated datasets. We applied standard Principal Component Analysis (PCA), kernel PCA and Autoencoder Neural Networks on 1360 datasets from four different measurement technologies. The latent feature spaces are tested for their ability to (a) reconstruct the original data and (b) improve predictive performance on validation datasets not used during the creation of the feature space. While linear techniques show better reconstruction performance, nonlinear approaches, particularly, neural-based models seem to be able to capture non-additive interaction effects, and thus enjoy stronger predictive capabilities. Our results show that low dimensional representations of the human transcriptome can be achieved by integrating hundreds of datasets, despite the limited sample size of each dataset and the biological / technological heterogeneity across studies. The created space is two to three orders of magnitude smaller compared to the raw data, offering the ability of capturing a large portion of the original data variability and eventually reducing computational time for downstream analyses.

## I. Introduction

Gene expression is the process by which genetic instructions control the various cell mechanisms through the synthesis of proteins and other gene products [1]. High-throughput technologies such as microarrays [2] and RNA sequencing [3] measure gene expression profiles in a fast and automated manner. Gene expression data are by nature high-dimensional, and, unfortunately, transcriptomics datasets typically have low sample size, due to either budget constraints or limited availability of samples (e.g., patients affected by a rare disease). These settings make the computational analysis of individual studies less accurate and statistically not robust [4]; furthermore, sophisticated algorithms may be required for addressing the intrinsic challenges that a large number of variables imposes.

Interestingly, many correlated variables exist in transcriptomics data, and thus there is redundant information which can be summarized by using, for example, dimensionality reduction techniques [5]. Dimensionality reduction transforms high-dimensional data into a reduced dimension representation with minimum loss of information [6]. Ideally, the dimensionality of the reduced representation should correspond to the intrinsic dimensionality of the data which is the minimum number of free parameters needed to account for the observed properties of the data. However, dimensionality reduction results might not be reproducible across different datasets due to the specificities of each study [7], [8], [9]. This also implies that a lower dimensional representation learned on a single study is almost certainly *not transferable* to other datasets, especially across different tissues, time points or experimental conditions.

In this study *we attempt to identify low-dimensional latent representations able to capture the biological information encoded in the whole human transcriptome across independent studies*. To this aim, we perform an extended analysis on a large compendium containing *1360 datasets* measuring various types of biological samples and conditions, such as healthy tissues, cancer subtypes, cell lines, etc., from three microarray and one Next Generation Sequencing RNA-seq platforms. A different low dimensional feature space is built for each platform. The constructed latent spaces are universal in the sense that heterogeneous datasets from a range of cell types, tissues and disease states are merged in a single large dataset for each platform. The ability of the latent features to preserve the original biological information is assessed on test datasets *never used during the derivation of the latent space*. For each test dataset, we quantify the ability of the latent representation (a) to reconstruct the original data and (b) to enhance the predictive modeling of phenotype information. Finally, in order to provide biological interpretation for the latent features, we compute their association with known biological pathways through Gene Set Enrichment Analysis (GSEA, [10]).

On all platforms we observe that the latent representations achieve better predictive performances than the original data, and allow a near-optimal reconstruction of the original datasets. Furthermore, latent features expected to carry more biological information are indeed associated with a larger number of pathways than less information-rich features. Thus, all together, our results provide evidence for the following, important points:

- building a global, low-dimensional representation for gene expression data that generalize on newly, unseen datasets is possible. This implies that gene expression data lie in a much lower manifold than their original raw data.
- low-dimensional representations whose information loss is minimal can ease the biological interpretation of transcriptomics data, by providing constructed features that can be correlated with known pathways and phenotypes.

The integration of gene expression datasets has been previously performed with two different approaches, namely late and early stage integration. The late stage integration is a ‘meta-analysis’ application, where each dataset is examined independently and the results are then combined together [11], [12], [13], [14]. In the early stage integration datasets are directly merged, eventually over common genes if measured on different microarray platforms [15], [16], [17], and the analysis is performed on the resulting super-datasets. Some of these works used an approach similar to ours: in [18], [19], about 5300 and 7100 samples from different studies were fused before applying PCA. The authors performed visual and cluster analysis and claimed that just the first few components had clear biological interpretation while the remaining contained irrelevant information. Interestingly, the study with the larger sample size concluded a larger intrinsic dimension of the latent space but the actual value remained unanswered. We identify three main methodological contributions:

- The utilization of tens of thousands of samples for the training of the dimensionality reduction methods. To put our contribution into perspective, the analysis on the same platform as in [18], [19] uses approximately 60000 samples resulting in an at least *eight-fold increase of the training data* relative to the previous studies. The laborious and time-consuming task of dataset collection is required in order to fully unveil the capacity of the explored dimensionality reduction methods.
- The use of *autoencoder neural networks* as a dimensionality reduction method for gene expression data.
- The employment of an automated machine learning pipeline for the assessment of the constructed latent feature space. We evaluate the dimensionality reduction methods based on their ability to predict the phenotypic information from the reduced datasets. Previous studies did not examine and quantify how those constructed latent feature spaces behave and generalize in terms of predictive performance on new unseen datasets.

## II. Methods

### A. Datasets

For our experiments we downloaded datasets, with at least 20 samples each, from BioDataome [20, http://dataome.mensxmachina.org/], a collection of *uniformly preprocessed* and *automatically annotated* datasets for data-driven biology. BioDataome hosts microarray gene expression datasets from the Gene Expression Omnibus database [21] and RNA-seq gene expression datasets from the recount database [22]. We downloaded 1185 microarray datasets of two different species (Homo Sapiens and Mus Musculus). For the Homo Sapiens datasets, 899 were measured with the Affymetrix Human Genome U133 Plus 2.0-**GPL570** and 86 with the Affymetrix Human Genome U133A-**GPL96** platform. The 200 Mus Musculus datasets were measured with Affymetrix Mouse Genome 430 2.0-**GPL1261**. We also downloaded 175 Homo Sapiens gene expression **RNA-seq** datasets. It is worth noting that all datasets belonging to the same platform measure exactly the same quantities, i.e., the same probesets for microarray data and the same gene count values for RNA-seq datasets.

In BioDataome, microarray data are preprocessed using the SCAN method [23]. SCAN processed and normalize each microarray profile *independently*, in contrast to other popular methods like Robust Multi-array Average (RMA, [24]) whose operation draws information from multiple samples at once. This feature ensures two desirable properties: (i) microarray data integration is not dependent upon the grouping of the individual sampling cross different datasets; (ii) performance assessment of predictive models on test sets is not hindered by dependencies artificially introduced during preprocessing. On the other hand, preprocessing methods for RNA-seq data customarily handle each sample in isolation, thus achieving the two properties above. Particularly, the recount2 pipeline [25] is used for all BioDataome RNA-seq datasets, further ensuring that all samples are normalized according to library size. Finally, BioDataome provides labels with disease terms from the Disease Ontology (http://disease-ontology.org/) for each dataset. The platforms and preprocessing methods used in this work do not give rise to missing data.

We manually labeled the samples of approximately 10% of the datasets for each platform with detailed phenotype information (GPL570: 80 studies, GPL96: 9, GPL1261: 20, RNA-seq: 18) which constitutes our test set. Phenotype labels correspond to information regarding disease states, cancer subtypes, smoking status, etc, as presented in Figure 1. Table I summarizes the statistical properties of the test sets with respect to their sample size for each platform.

**Fig. 1:**
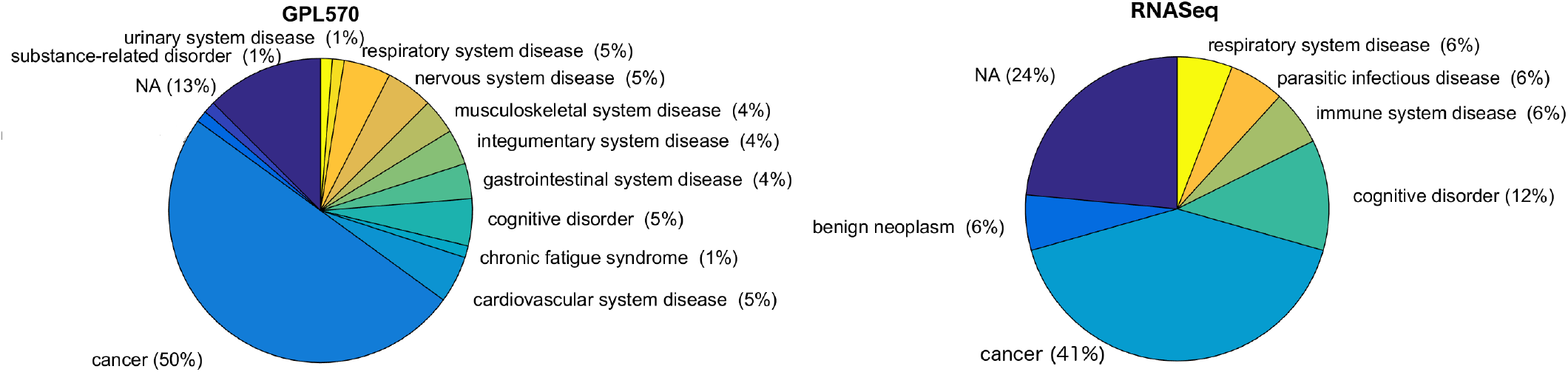
Pie charts with the disease/state categories of the test sets for the four platforms. Approximately half of the human gene expression datasets study cancerous samples.

**TABLE I:** Statistical information about the datasets for each platform. Dimension corresponds to the number of probe-sets.

| Platform | Dimension | No. test sets | Mean | Std | Min | Max |
| --- | --- | --- | --- | --- | --- | --- |
| GPL570 | 54675 | 80 | 107.20 | 54.49 | 25 | 200 |
| GPL96 | 22834 | 9 | 55.11 | 26.39 | 20 | 100 |
| GPL1261 | 45101 | 20 | 49.65 | 36.93 | 20 | 144 |
| RNA-seq | 23779 | 18 | 239.56 | 145.36 | 61 | 520 |

The remaining datasets were merged creating 4 different super-datasets comprising 59864, 3331, 7282 and 21609 samples from GPL570, GPL96, GPL1261 and the RNA-seq platform, respectively. Note that the sample size is either less or approximately the same as the dimension (i.e., the probe-set size) of the samples in all platforms.

### B. Experimental Protocol

Our dimensionality reduction analyses follow the protocol depicted in Figure 2. In brief, in each analysis we merge the transcriptome profiles from several datasets, creating large super-datasets containing thousands of samples each. We then constructed several low dimensional feature spaces for each super-dataset by applying the following dimensionality reduction methods: PCA [26], kernel PCA [27] and the state-of-the-art Autoencoder Neural Networks [28]. We tested the ability of the constructed latent representations to preserve the original biological information by assessing their capability of enhancing predictive modeling and reducing the reconstruction error on newly-seen datasets, i.e., datasets not included in the training of the dimensionality reduction methods. We examined and evaluated various compression magnitudes in order to identify the lowest latent feature dimension capable of outperforming the predictions obtained on the full datasets.

**Fig. 2:**
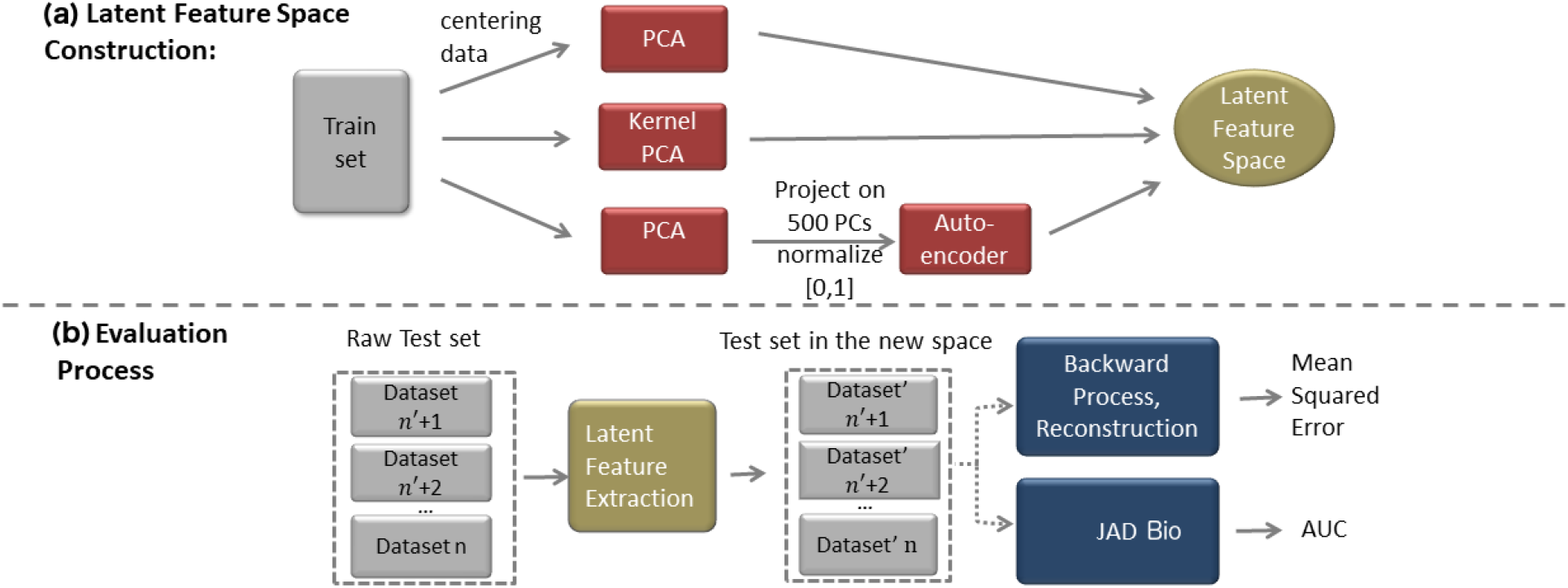
Outline of our analysis pipeline: (a) Dimensionality reduction is applied using several linear and nonlinear methods. We constructed latent feature spaces with variable dimension values. For the Autoencoder, we first projected to the 500 strongest PCs in order to reduce the number of neural network’s parameters. (b) The remaining 10% of the studies (i.e., the Test set) are projected onto the constructed latent feature space. Then, we evaluated the quality of the dimensionality reduction algorithms in terms of both reconstruction error in the original space and classification performance using JAD Bio software, a fully-automated machine learning pipeline.

The first step of the dataset integration consists in simply concatenating a number of different datasets, an operation possible thanks to datasets from the same platform measuring exactly the same quantities, i.e., the same probesets for microarray data and the same gene count values for RNA-seq datasets. Particularly, for each platform, we merged 90% of the collected studies and generated an integrated dataset with thousands of samples which served as the training set to the dimensionality reduction algorithms. Fig. 2(a) depicts the latent feature construction processes. PCA as well as kernel PCA were applied on the original dimension. However, the training of an Autoencoder neural network was not only impractical but also ill-posed on the original, high-dimensional data, due to the fact that the number of Autoencoder’s parameters was two to three orders of magnitude larger than the number of samples making the training infeasible and prone to overfitting. In order to overcome this issue, we performed an initial dimensionality reduction using PCA and kept the 500 largest Principal Components (PCs) of each set. It is worth noting that 500 PCs suffice to explain 96% of the (standardized) data variance on average (i.e., one minus the ratio between reconstruction error and squared Euclidean norm). Hence, without loss of any significant information, these 500 PCs were normalized to take values in the interval [0, 1] and then fed as input to the Autoencoder for further dimensionality reduction.

The evaluation of the dimensionality reduction performance is measured with the reconstruction error using the hold-out datasets. Specifically, we held out 10% of datasets on which to estimate the reconstruction error, defined as the mean squared error between the original and the reconstructed datasets. We additionally employed the JAD Bio software to measure the phenotype prediction performance of the projected datasets as shown in Fig. 2(c). As an evaluation metric, we reported the Area Under the ROC Curve (AUC). AUC is a reliable metric since it is invariant of the sample size of each category. The AUC on the original datasets was also computed and used as a reference.

### C. Dimensionality Reduction Methods

#### 1) Principal Component Analysis (PCA)

The most popular technique for dimensionality reduction is PCA [26]. It performs an orthogonal transformation converting a set of possibly correlated features into a smaller set of linearly uncorrelated variables. The new coordinate system projects the data into the directions with the highest variance. In mathematical terms, PCA attempts to find a linear mapping *M* that maximizes the cost function *trace*(*M*^**⊤**^*cov*(*X*)*M*), where *cov*(*X*) is the sample covariance matrix of the data matrix *X*. It turns out that this is done by finding the eigenvalues and eigenvectors of the sample covariance matrix, i.e., the PCWs. An efficient method for estimating the principal components is through Singular Value Decomposition (SVD). SVD factorizes the data matrix as *X* = *USV^T^* where the right singular vectors *V* are equal to the eigenvectors of the sample covariance matrix, hence, equal to the PCWs.

#### 2) Kernel PCA

Kernel PCA [27] generalizes standard PCA to nonlinear spaces. The idea behind kernel PCA, is to initially use a nonlinear transformation function *ϕ*(*x*) from the original feature space to a nonlinear and larger feature space and then perform PCA. The transformation map *ϕ*(*x*) is typically extremely high-dimensional making the mapping inefficient. The explicit calculation of the new feature space can be avoided using the kernel trick. The kernel trick refers to creating a kernel function (similarity function) which is a function over pairs of data points in raw representation *K*(*x,x′*) = *ϕ*(*x*)^**⊤**^*ϕ*(*x′*). The kernel function of a dataset with *N* samples is a *N × N* matrix whose eigenvectors are equivalent to the principal components in the latent space created by *ϕ*(*x*). The most widespread kernels are Polynomial and Gaussian (a.k.a., radial basis functions; RBFs). We apply Polynomial kernels of degree 2 and 3 and Gaussian kernel with hyper-parameter set to one over the dimension of the samples. Preliminary experiments on hyper-parameter tuning for Gaussian kernel showed that our choice is on average close to the optimal value. Finally, we note that both Kernel PCA and standard PCA are limited to produce up to n-1 components, where *n* is the minimum between the number of samples and number of measurements.

#### 3) Autoencoder Neural Networks

Hinton and Salaktutdinov [28] proposed Deep Learning Autoencoders in order to convert high-dimensional images to low-dimensional codes with compact information. Autoencoder is a special type of neural network that has two parts: the encoder, *f*, that maps raw input *x* into a nonlinear representation *h* = *f*(*x*) and the decoder, *g*, that maps h back to the original space. Neural networks are able to capture multi-modal aspects of the input distribution and, hence, capture nonlinear interactions between the original features. Additionally, the expressiveness of a neural network is enhanced with the use of more hidden layers allowing them to compactly represent highly nonlinear functions. Despite their success in image and speech recognition, only recently neural networks have been used in biological research [29], [30], [31] primarily due to the limited number of samples for training. Indeed, neural networks require large datasets and computational power for satisfactory learning without overfitting. In this paper, the issue of limited samples is alleviated by dataset integration which results in tens of thousands of gene expression samples.

In neural network modeling, there are several hyperparameters that should be tuned a priori before the learning phase, the most significant being the structure of the network. The learning rate, the number of epochs and the size of batches in stochastic gradient descent are additional hyper-parameters that need to be specified. We performed a series of initial analysis on a limited set of data for determining suitable values for all hyper-parameters. We decided to use three layers both for the encoder and for the decoder with the number of units for the intermediate layers was 700 and 500. The sigmoid function was chosen as the transfer function for both hidden and output units while the Autoencoder was pre-trained using a greedy layer-wise unsupervised learning algorithm which employs Restricted Boltzmann Machines [32]. Since the input data are normalized to lay in the interval [0, 1], they can be thought as probabilities or intensities. Therefore, the crossentropy loss function between the input and output intensities, which does not require any Gaussianity assumption for the error, was used as the objective function to be minimized [33], [28]. Finally, we set the batch size to 200 while the learning rate was 0.1 for the first 1K epochs, 0.03 for the next 1K and 0.01 for the last 1K epochs.

### D. Predictive Modeling

The evaluation based on predictive analytics is performed on 127 annotated datasets, a.k.a., the test sets which corresponds roughly to 10% of the available datasets (details can be found in supplementary files 1–4). For each of these datasets the transcriptomics measurements were used as predictors, while phenotype information was used for manually creating an outcome coherent with the specific design of each study, categorizing samples in known groups (e.g., disease vs. healthy controls).

Predictive modeling is carried out with Just Add Data (JAD) Bio software tool (www.jadbio.com) which produces a conservative estimation of AUC. JAD Bio implements a fully-automated machine learning pipeline for creating a predictive model given a training dataset, and an estimate of its predictive performance. JAD Bio first selects the algorithms for transformations, imputation methods for missing values, feature selection, and classification to apply; it also selects the performance estimation protocols. The choices depend on the size and type of the data and the user preferences. Once a set of suitable algorithms is chosen, JAD Bio tries several configurations, i.e., combinations of algorithms and values of their hyper-parameters, in an effort to identify the configuration that optimizes predictive performance. Notably, JAD Bio has already been used in several high-level publications [34], [35], [36].

In terms of algorithms, JAD Bio trains several basic and advanced, linear and non-linear, multivariate machine learning and statistical models. For each algorithm, it tests different values of their hyper-parameters using a grid-search in the space of hyper-parameters. For the sample size and dimensionality of the datasets included in our experimentation, JAD Bio adopted the following solutions:

- The configurations space is explored through a grid search, while repeated, stratified cross-validation [37] is used for determining the best configuration. The number of folds is set to the minimum between 10 and the number of elements in the rarest class. The best configuration is then used to produce the final models on all data.
- The Statistically Equivalent Signatures (SES) algorithm [38] is used for feature selection, in order to build parsimonious, yet maximally predictive, models. SES hyperparameters *α* and *maxK* are optimized within values {0.01,0.05,0.1} and {2,3,4}, respectively.
- Decision Trees, Random Forests, Support Vector Machines with Gaussian and Polynomial kernels, and Ridge Logistic Regression are used as modelers. JAD Bio tests the cost for all the different kernels of the SVMs with values {0.001, 0.01, 0.1, 1, 10, 100}, the gamma for the polynomial and gaussian kernel SVMs with {0.001, 0.01, 0.1, 1, 10, 100}, and the degrees of the polynomial kernel SVMs with {2, 3, 4}. The penalty parameter lambda of the Ridge Logistic Regression is tested with the values {0.001, 0.01, 0.1, 1, 10, 100}. For Random Forests, JAD Bio trains 1000 trees, with minimum leaf sizes (1, 2, 3, 4, 5), and tests different values of the number of variables to select at random for each decision split using the formula 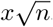, where *x* takes the values {0.33, 0.66, 1, 1.33, 1.66, 2} and *n* is the number of the variables that the dataset contains.

The cross-validated performance of the best configuration is known to be optimistic due to the multiple tries [37]; JAD Bio estimates and removes the optimism using a bootstrap method before returning the final performance estimate [39]. The final performance estimates of JAD Bio are actually slightly conservative. Finally, no feature transformation or imputation of missing values was required for the data included in our analysis.

### E. Gene Set Enrichment Analysis

We are interested in understanding the biological relevance of the constructed features. To this aim, we performed Gene Set Enrichment Analysis (GSEA) [40] on the human microarray data using principal component weights (PCWs) as enrichment scores, which correspond to the columns of matrix V in PCA. Gene sets are predefined groups of genes that are involved in the same biological process (pathway). A popular pathway database is the Kyoto Encyclopedia of Gene and Genomes (KEGG, [10]) from which we downloaded 186 different gene sets. For each gene set (A), we split each of the 200 strongest PCWs into the ones included in (A), denoted as A1 and the rest, denoted as A2. Intuitively if A1 is statistically significantly different from A2 we say that this gene set is being enriched. We used the Wilcoxon Rank Test [41] as statistical test and False Discovery Rate (FDR) level 0.05 [42]. For robustness purposes, we do not take into account probesets that are not associated to any gene as well as probe-sets that point to several genes. Finally, we exclude gene-sets that belong to KEGG but have less than 10 genes measured by the analyzed platforms. The remaining gene-sets are 143 and 161 for the GPL570 and GPL96 platforms, respectively.

## III. Results

Figure 3 presents the mean reconstruction error (upper row of panels) as well as the mean AUC (lower row of panels) evaluated on the newly-seen datasets. We chose to compute the performance metrics at a wide range of dimension values for the latent feature space and produce a performance profile for each method and each platform. As expected, the reconstruction error is increased as the dimension of the latent space is decreased. The Autoencoder (green lines) either had the lowest reconstruction error or was in par with PCA (blue lines) for the platforms with large sample size (i.e., GPL570 and RNA-seq). For GPL570 platform, for instance, 74% of the relative variance is kept using Autoencoder which is 10% higher than PCA’s relative variance when the dimension of the latent feature space is 20. In contrast, PCA achieves better reconstruction results when the total sample size is small (i.e., GLP96 and GPL1261). The nonlinear Kernel PCA (cyan, red and black lines) produced relatively higher reconstruction error in all platforms. However, we would like to remark that the reconstruction in Kernel PCA requires taking the preimage of the kernel features back in the original space. This process is not always feasible and we have used approximation algorithms [43] whose error adds up to the reconstruction error.

**Fig. 3:**
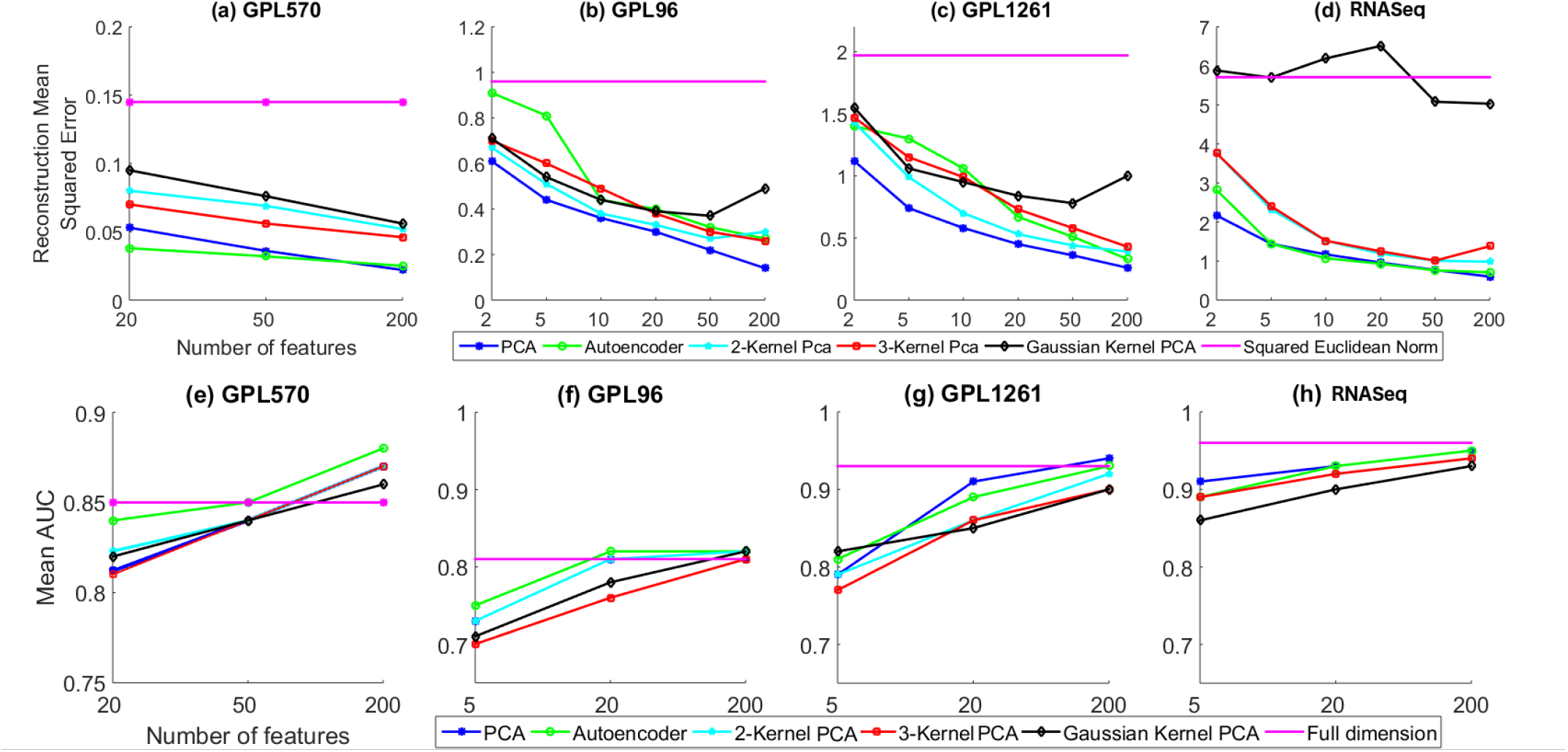
Performance assessment of dimensionality reduction techniques on test datasets for each platform in “within platform” integration scheme both in terms of reconstruction error (upper row of panels) and classification through mean AUC (lower row of panels). It is evident that the reconstruction error decreases with the number of latent features. The reference point (pink line) is the squared Euclidean norm of the sets and it is practically the variance. PCA (blue line) and Autoencoder (green line) are the dominant methods in terms of reconstruction accuracy. Regarding classification performance, it is verified that a strong reduction to 20 dimensions or less, results in a loss of predictive accuracy. Nevertheless, with a latent feature space of 200 dimensions, the prediction performance is equal or better compared to the performance of the original data for all platforms.

The predictive performance as measured by mean AUC was higher for the Autoencoder method than for PCA, particularly using Homo Sapiens microarray datasets. On the other hand, Kernel PCA had a consistently modest but slightly worse performance on all platforms. For GPL570 (Figure 3(e)), we were able to get comparable results with the reference mean AUC (pink line) when Autoencoder (green line) was applied for the construction of a 50-dimensional latent feature representation. Moreover, the performance is improved by 3% compared with the reference mean AUC when the latent feature dimension is set to 200. Actually, twelve datasets had increased AUC by at least 10% while only one dataset’s performance deteriorated by the same percentage compared to the original dataset’s performance.

For GPL96 (Figure 3(f)), Autoencoder and PCA with Gaussian and 2-polynomial kernels improved the classification accuracy compared to the reference, again, for 200 latent space dimensions. Actually, all dimensionality reduction methods returned equal or slightly higher results than the original highdimensional data when the latent feature space dimension is 200. Interestingly, Autoencoder, PCA and 2-kernel PCA for GLP96 were able to get better or equal classification performance than the reference even for 20 dimensions showing that the gene expression data might be represented with a small number of compressed features. The other two platforms (Figure 3(g) and Figure 3(h)) showed similar qualitative behavior. The results for these two platforms were slightly worse because the reference mean AUC (pink lines) is about 95% which is close to perfect classification. Finally, we report the variability of the obtained results. The average standard deviation of the mean AUC is 0.023 when the original data are used and 0.018 & 0.19 when the latent features with 200 dimensions were used for the Autoencoder and the PCA, respectively.

The above results indicate that gene expression data are indeed redundant with two to three orders of magnitude lower intrinsic dimensionality. Nevertheless, in order to preserve the predictive performance, gene expression data can be reduced to a latent feature space with approximately 200 dimensions. Additionally, the size of the training set crucially affects the performance of the dimensionality reduction methods. The intrinsic interactions between genes and/or experimental conditions can be more effectively captured by applying sophisticated reduction methods such as the Autoencoder given the availability of a large number of samples. To support this claim we conduct an experiment where the dimension of the latent representation is kept fix while the size of the train set it varied. Figure 4 shows the Autoencoder’s improvement in terms of both the reconstruction error and the mean AUC for 20 (red) and 200 (blue) latent dimensions as a function of the training sample size. The reconstruction error is a decreasing function of the training sample size while the mean AUC is most of the times an increasing function showing that the Autoencoder is crucially benefited from the increase in sample size. Finally, we can also conjecture that supplementing the train set with additional data, the performance of Autoencoder as a dimensionality reduction method will be further improved.

**Fig. 4:**
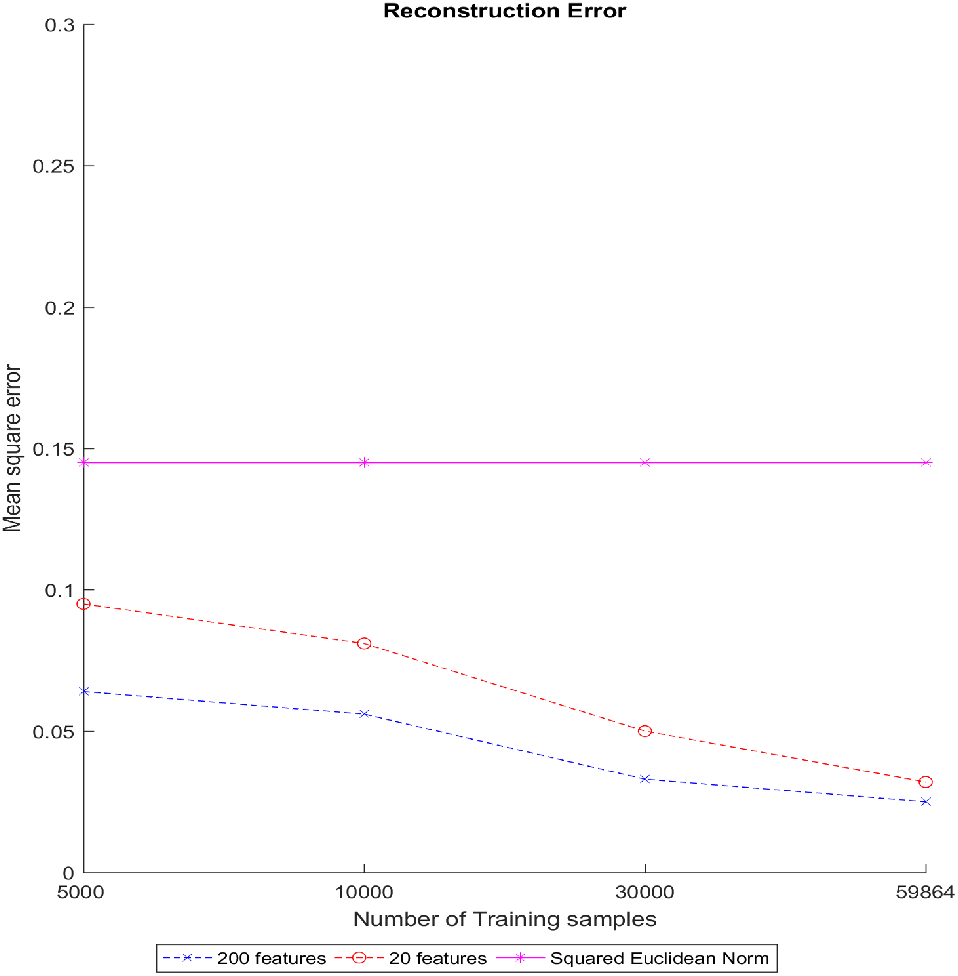
The reconstruction error (left panel) and the mean AUC (right panel) as a function of the train set size when Autoencoder with 20 (red) and 200 (blue) latent space dimensions is utilized. Both metrics improve as the sample size is increased.

### A. Gene Set Enrichment Analysis

Figure 5 reports the GSEA FDR adjusted p-value for each gene-set on every PCW from platform GPL570. A similar figure is reported in the additional file 3 for the GPL96 platform. The x-axis represents gene-sxets while the y-axis corresponds to PCWs. Red colored cells correspond to highly enriched pathways (adjusted p-value close to zero), while a blue cell corresponds to a non-significant pathways (adjusted p-value close to one). We used hierarchical clustering with Euclidean distance and complete linkage for grouping both PCWs and pathways, and we identified the optimal number of clusters by maximizing the Ratkowsky–Lance criterion [44]. The results can be summarized as follow:

- All pathway are found enriched for at least one PCW, with the only exception of 1 and 5 never-enriched pathways in GPL570 and GPL96, respectively. This indicates that the computed PCWs carry information related to most of known biological mechanisms.
- The strongest PCWs enrich more pathways than the subsequent latent features, demonstrating that these PCWs have most of the crucial biological information. In Figure 5, PCWs are clustered in three different groups, roughly following the PCWs’ order: the first cluster, in red in the rows’ dendrogram, contains the first 9 PCWs along with L14-L16, L18, L24 and L34. Each PCW from this first cluster enriches 44.7 pathways on average. The second rows’ cluster (green) mainly contains PCWs from L20 to L70, which enrich on average 8 pathways each, while the third rows’ cluster (blue) contains the remaining latent features and has the lowest number of enriched pathways on average (2.7). A similar behavior can be observed for the GPL96 platform (additional file 5).
- Seven pathways are enriched in many of the latent features for both platforms: Systemic Lupus Erythematosus, Ribosome, Complement and Coagulation Cascades, Alzheimer Disease, Parkinson Disease, Huntington Disease and Oxidative Phosphorylation. The latter four pathways also share a large proportion of their genes: each pair among them has a Jaccard coefficient over 0.4, where the Jaccard coefficient among two sets is defined as the ratio between the intersection and the union of their elements. In Figure 5 the seven pathways correspond to the leftmost columns, which are clustered together in the red and blue columns’ clusters.

**Fig. 5:**
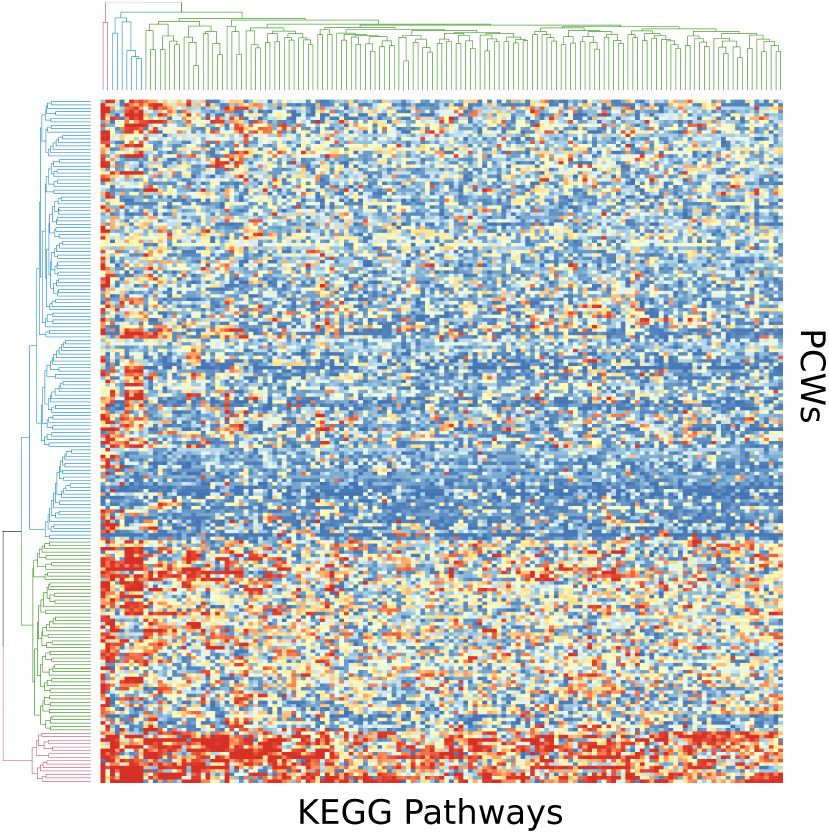
Heatmap representing the adjusted p-values for the enrichment analysis. Each cell at position (*i, j*) indicates the adjusted p-value for the *i*-th latent feature and *j*-th gene-set. Red color indicates high significance (adj. p-value close to 0), while blue color the opposite.

### B. Batch effects

In all our analyses we do not take into account batch effects between datasets or between platforms, letting the dimensionality reduction methods implicitly handle them. Figure 6 shows the first PCs for the never-seen datasets obtained through PCA analysis of the integrated Homo Sapiens microarray platforms. The left panel of Figure 6 presents the two strongest PCs which almost perfectly separate the test datasets of the two platforms. In the right panel, we show the 3rd, 4th and 5th strongest PCs for 10 test datasets from the GPL570 platform. It is evident from both plots that between-datasets batch effects are present; however, our results suggest that they do not affect the prediction performance. Hence, the PCs convey simultaneous, multi-source information for (a) both the experimental and the disease conditions of the samples and (b) biological pathways (see the above subsection). In essence, we efficiently learn the combined global map of the genome and the global map of the measurement characteristics of the various technologies and laboratories. It is also noteworthy that in order to learn both the genome and the experimental maps the dimension of the latent representation should be large enough. In a dedicated experiment (not shown here), we compare the reconstruction error with and without batch effect removal^1^ and observe that a substantial difference when the dimension is below 50. This is reasonable because when the dimension is small, the overall sample variability cannot be fully captured by a low-dimensional latent space. However, when the dimension is above 100 then batch effect removal did not offer any performance edge to the dimensionality reduction approaches (at least the linear ones).

**Fig. 6:**
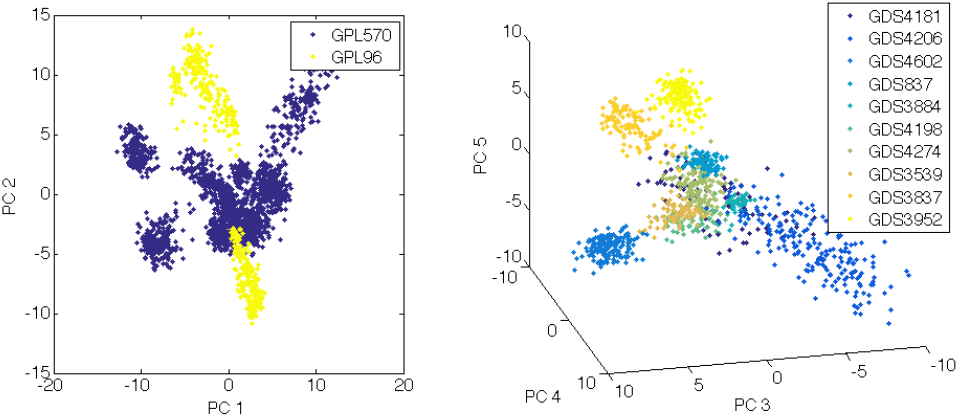
The projection of unseen datasets from both Homo Sapiens microarrays on their strongest PCs. The left panel shows all sets that have been kept out of the training procedure projected on the two strongest PCs. The datasets from GPL570 (dark blue) and GPL96 (yellow) are almost completely separated. The 3rd, 4th and 5th strongest PCs are able to distinguish each dataset as the right panel reveals. The ten unseen datasets of GPL570 platform are visually separated forming distinct clouds of data points.

## IV. Discussion

Gene expression datasets have both low sample size and high dimension making data integration as well as dimensionality reduction two mandatory procedures for their robust and reliable statistical and computational analysis. In this paper, we integrated hundreds of studies from four different platforms and two different technologies, and extensively investigated the construction of latent feature spaces using three different dimensionality reduction methods (PCA, Kernel PCA, Autoencoder Neural Network). Despite not being the first who applied dimensionality reduction techniques on gene expression data, our novelty stems from the facts that (a) we are utilizing hundreds of datasets that belong to the same or different platforms, (b) we extensively search for the appropriate dimension of the latent representations, and (c) we test the constructed latent feature spaces on newly-seen datasets.

We demonstrated that a large dimensionality reduction is possible without affecting the underlying biological information. Particularly, we showed that the mean prediction performance on new datasets is markedly improved in the constructed latent feature space compared to the raw features. The Autoencoder method demonstrated the best performance in terms of classification accuracy, especially when the number of available samples is large, showing that handling nonlinear interactions of genes allows to achieve better classification performance. Overall, the improvement in performance indicates that the new features not only summarize the information contained in the original measurements, but expose such information in a way that is usable in a more effective way by the machine learning methods that we subsequently applied.

In our analyses the optimal size for the latent representation is approximately 200 dimensions. This value is larger than the reported in previous studies [18] indicating that there is crucial biological information in higher dimensions that boosts the machine learning algorithms to achieve better predictive results. The selection of the optimal dimension for the latent space requires a criterion that balance between the size/complexity of the latent representation and the predictive performance of the reduced datasets. For instance, if the interest is mainly in the accuracy, the latent dimension can be increased until the mean AUC is only marginally improved.

The constructed latent features are also informative with respect to known biological pathways, with the strongest PCWs enriching a larger number of pathways compared to the weaker ones. These results are not trivial: the construction of the latent features and the definition of the biological pathways are two totally separated procedures. The presence of enrichment suggests that the dimensionality reduction is able to encode actual biological mechanism in the latent features. A deeper understanding regarding whether the characteristics of the reduced features could be dictated by the underlying biological processes would require further study.

Finally, we point out that dimensionality reduction also leads to significant gains in computational time for an extended predictive analysis. A full-scale predictive analysis using the automated machine learning tool JAD Bio on gene expression data requires from 5 to 30 minutes for completion while the same analysis on the reduced latent feature space takes from 10 seconds to 2 minutes. The variability is explained by the different sample sizes as well as by the difficulty of the classification task for each dataset. Dimensionality reduction methods have the additional advantage of naturally taking care of gene overrepresentation in microarray datasets, i.e., the presence of multiple probesets for the same gene, which creates groups of collinear measurements. Such groups are elegantly projected into the same components.

## V. Conclusions

The collection of a broad number of microarray and RNA-seq datasets enabled us to perform a large-scale analysis and assessment on three different dimensionality reduction techniques. We showed that the within platform integration of datasets when they are uniformly preprocessed resulted in the construction of low-dimensional latent representations that were able to increase the average performance of predictive modeling with the neural network Autoencoder producing the best results. Finally, the constructed latent features are also informative with respect to known biological pathways, with the strongest PCWs enriching a larger number of pathways compared to the weaker ones. The presence of enrichment suggests that the dimensionality reduction is able to encode actual biological mechanism in the latent features.

## Footnotes

1 We defined as a batch effect the statistical mean value of each dataset.

